# PlotXpress, a webtool for normalization and visualization of reporter expression data

**DOI:** 10.1101/2021.07.08.451595

**Authors:** Elias Brandorff, Marc Galland, Joachim Goedhart

## Abstract

In molecular cell biology, reporter assays are frequently used to investigate gene expression levels. Reporter assays employ a gene that encodes a light-emitting protein, of which the luminescence is quantified as a proxy of gene expression. Commercial parties provide reporter assay kits that include protocols and specialized detection machinery. However, downstream analysis of the output data and their presentation are not standardized. We have developed plotXpress to fill this gap, providing a free, open-source platform for the semi-automated analysis and standardized visualisation of experimental gene reporter data. Users can upload raw luminescence data acquired from a reporter gene assay with an internal control. In plotXpress, the data is corrected for sample variation with the internal control and the average for each condition is calculated. When a reference condition is selected the fold change is calculated for all other conditions, based on the selected reference. The results are shown as dot plots with a statistical summary, which can be adjusted to create publication-grade plots without requiring coding skills. Altogether, plotXpress is an open-source, low-threshold, web-based tool, that promotes a standardized and reproducible analysis while providing an appealing visualization of reporter data. The webtool can be accessed at: https://huygens.science.uva.nl/PlotXpress/

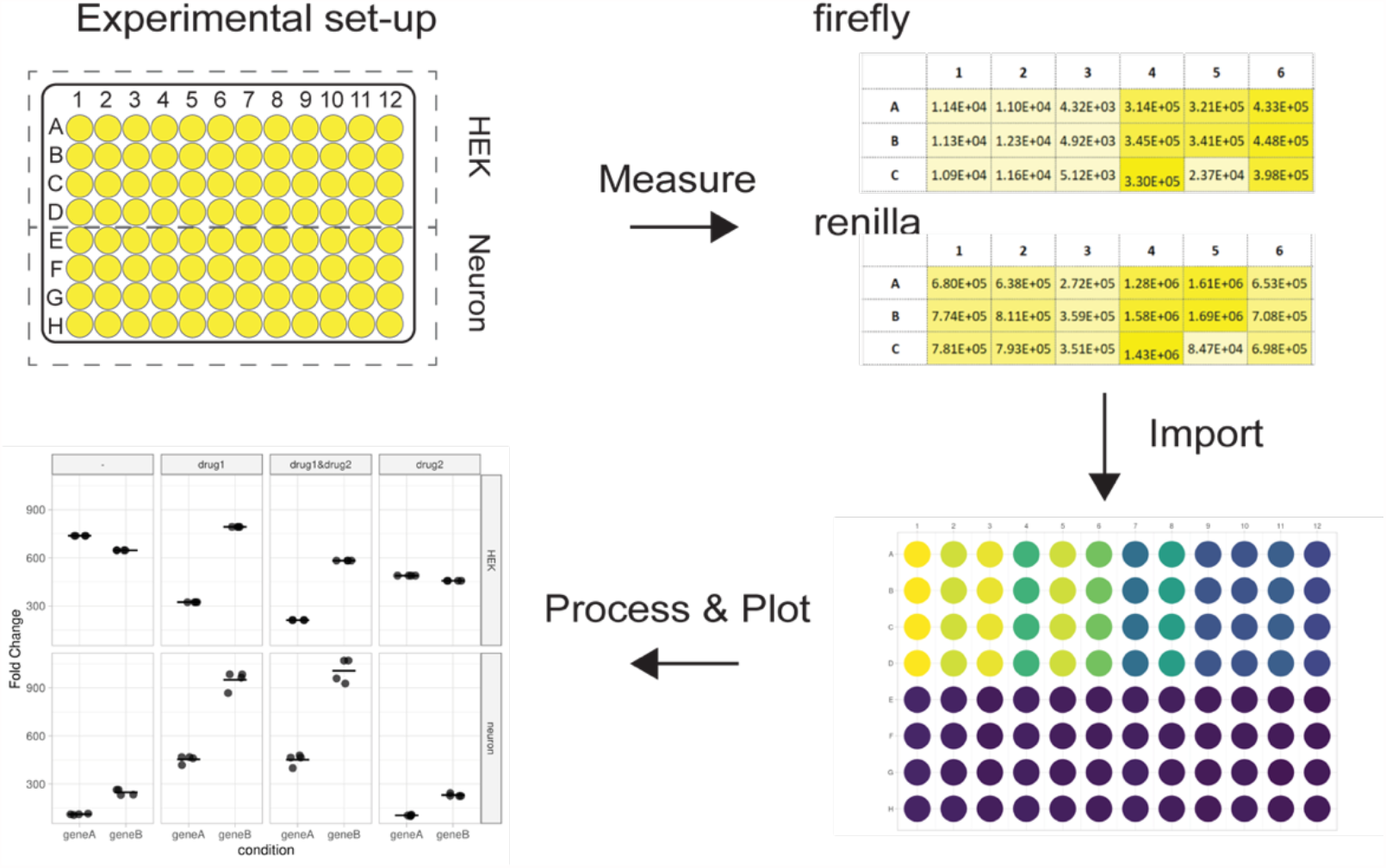

## Introduction

Reporter gene assays are popular tools to investigate gene expression dynamics in cell biology (Barriscale et al., 2014; Liu et al., 2009; Schenborn and Groskreutz, 1999). In most systems, the reporter is a gene coding for a protein that emits light when it binds a substrate. An example is the firefly (*Photinus pyralis*) luciferase that emits light when it binds the substrate luciferin (Himes and Shannon, 2000). The luminescent light can be detected with dedicated equipment and reflects the expression of the reporter gene.

The expression level of the reporter gene is used as a proxy of the capacity of a DNA sequence to regulate gene expression under various experimental conditions. To this end, the DNA sequence of interest is cloned upstream of the reporter gene in a vector which is subsequently transfected into cells. Reporter gene activity can be compared between variations of a regulatory DNA sequence, for example by removing a transcription factor binding site or by modulating tandem repeat expansions in a promoter sequence (Rodriguez et al., 2020; Moparthi and Koch, 2020). Another option is to measure the effects on reporter activity of treatments such as pharmaceutical compounds or overexpression of transcription factors that interact with the sequence of interest (Asamitsu et al., 2021). In addition, reporter assays have been used to study the relation between microRNAs and cancer progression (Wang et al., 2021; Chengling et al., 2021). In the so-called “Dual-Luciferase® Reporter Assay System” (Promega: https://nld.promega.com/resources/pubhub/in-vivo-evaluation-of-regulatory-sequences-by-analysis-of-luciferase-expression/), an internal control is used to correct for variations in cell density and transfection efficiency. This internal control is co-transfected and consists of a vector with a constitutively active promoter that drives expression of another luciferase. This second luciferase emits a different color of light and is, in our example, derived from *Renilla reniformis* (sea pansy). Transfection of the internal control is kept the same in each well and reflects transfection efficiency and cellular protein production. A given sequence of interest is cloned into a luciferase expressing plasmid in order to investigate its activity as a gene promoter. The transfection of an empty vector control, without this sequence, is an important reference condition and required to correct for unintended effects of the plasmid alone. For example, in some vectors, luciferase is expressed under control of an SV40 promoter element. A transcription factor binding site (TFBS), cloned upstream of this promoter will influence its activity and the expression of luciferase. The measured effect is a composite of promoter activity *plus* the TFBS. The empty vector control allows normalization for baseline promoter activity and isolation of the influence of the TFBS alone. Here, we will call the empty vector condition the *reference* condition.

In reporter assays with multiple reporter constructs, cell types, treatments and reference conditions, experiments may increase rapidly in size and complexity, leading to challenging downstream analyses (Schagat et al. 2007). There is currently neither standardized method nor computational tool for the processing of reporter assay data and its visualization. PlotXpress was designed to simplify the analysis process of complex reporter assays by providing an online tool with a standard for data processing and visualization. PlotXpress was built following the philosophy of transparent and state-of-the-art data visualization implemented in the data visualization app PlotsOfData (Postma & Goedhart, 2019). In addition to a streamlined analysis, plotXpress enables transparent communication of the data. Instead of only providing averaged data and error bars to summarize gene reporter data in bar graphs, plotXpress produces dot plots maintaining individual data points. Data in both wide and tidy format (Wickham, 2014) can be provided. As coding skills are not required, plotXpress is a readily available low entry-level application that democratizes the processing and visualization of dual reporter expression data.

### Availability & code

The plotXpress web tool can be accessed at https://huygens.science.uva.nl/PlotXpress/ (and a mirror is available at: https://goedhart.shinyapps.io/PlotXpress). Example data is provided to explore the functions of the app. The plotXpress code is written in the R programming language (https://www.r-project.org) using the following packages: ggplot2 (Wickham, 2016), tidyr, magrittr, readr, stringr, dplyr and readxl which are all part of the tidyverse suite of packages version 1.3.1 (Wickham et al., 2019), Shiny (https://CRAN.R-project.org/package=shiny), and DT (https://CRAN.R-project.org/package=DT).

This manuscript documents version 1.0.0 of the webtool which is archived (together with the example data) at Zenodo: https://doi.org/10.5281/zenodo.4980068

Background information, updates of the code and version releases will be published on GitHub: https://github.com/ScienceParkStudyGroup/PlotXpress. GitHub is the preferred channel for communication regarding issues and feature requests.

### Example Data

The plotXpress app comes with an example dataset and a file that describes the experimental conditions for each well ‘Tidy_design.csv’ (**Fig 1**). Both files, ‘DualLuc_example_data.xlsx’ and ‘Tidy_design.csv’, are also available online at https://github.com/ScienceParkStudyGroup/PlotXpress.

**Figure 1.**
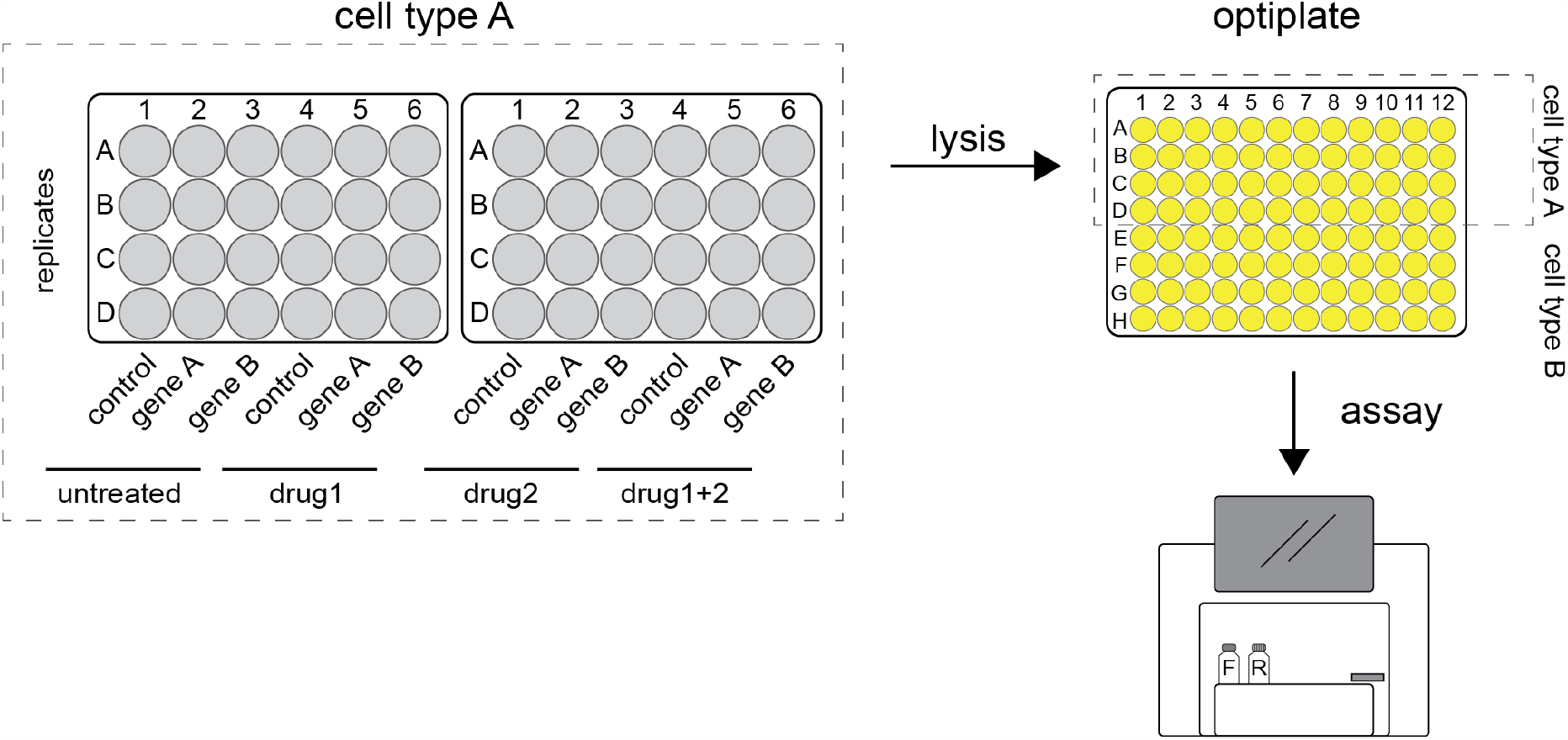
Graphic representation of a typical reporter assay. Firefly and renilla expressing plasmids are transfected into cells grown in 24-well plates. After incubation the cells are harvested and lysates pipetted into 96-wells optiplates. These are loaded into a plate reader where the luciferase substrates are added. Once the reactions are initiated, luminescence is detected, and readings are stored digitally.

### Data upload

Users can upload their data in two different ways. The first option is for data acquired with the Promega GloMax plate reader. The alternative is a general-purpose option that accepts data in a tidy format.

#### GloMax data upload

The output of the GloMax plate reader (Promega) is a spreadsheet XLSX format with two tables containing firefly and renilla luminescence signals stored in a 96-well lay-out. PlotXpress reads the cells with the firefly and renilla readings and provides a graphic overview of the experiment by showing a 96-well plate where signal intensity is colour coded (**Fig 2**).

**Figure 2.**
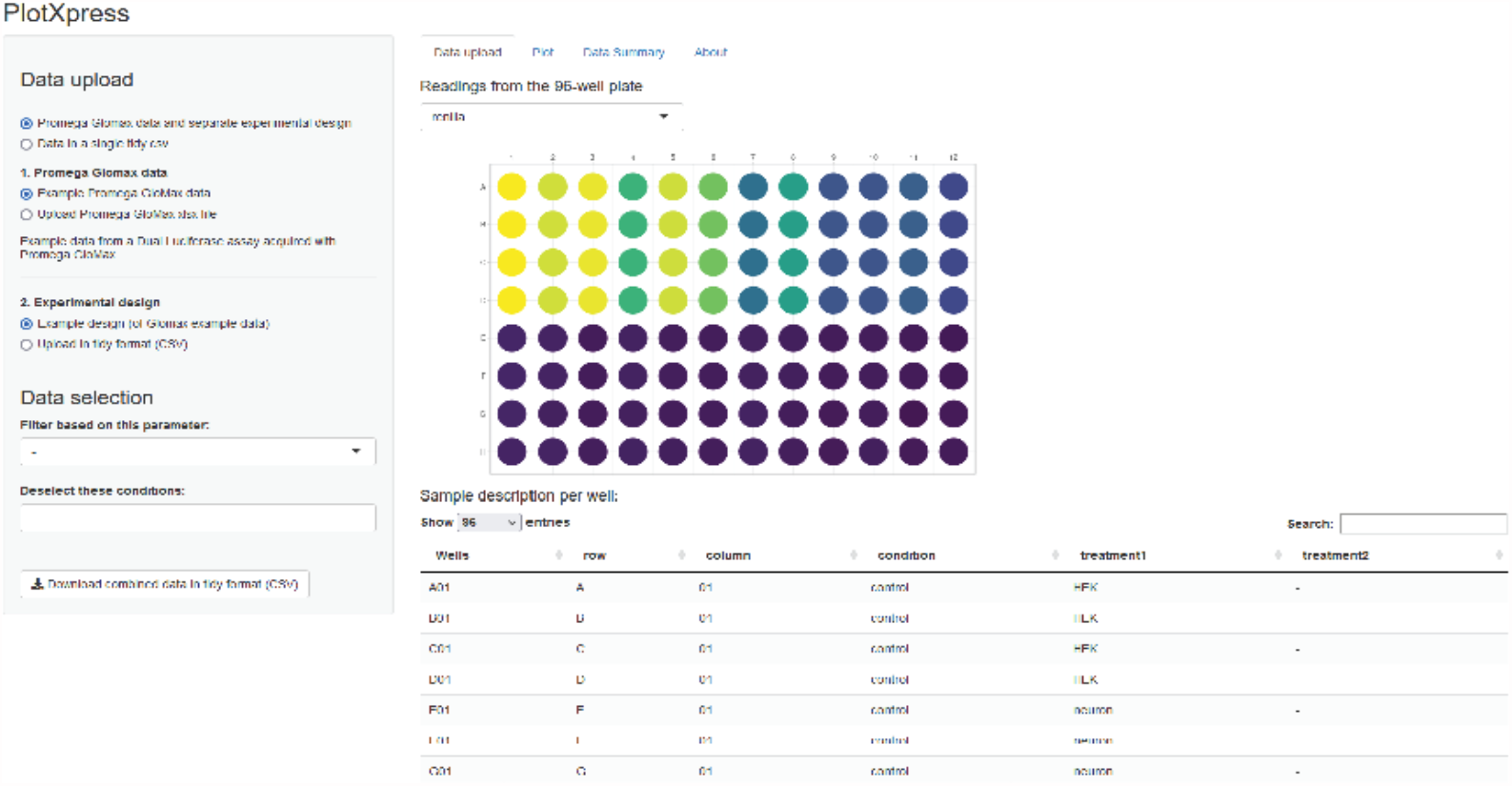
PlotXpress data upload. Screenshot of plotXpress app showing a 96-well format with signal intensities represented by false colour.

A separate table in CSV format with the experimental conditions per well is required for the data processing. A template is available for download within the app or online: https://github.com/ScienceParkStudyGroup/PlotXpress/blob/master/Tidy_design.csv

The firefly and renilla reads from the GloMax data are converted into a tidy format and merged with experimental conditions that are taken from the uploaded design (**Fig 3**). The resulting tidy dataframe is used for data processing.

**Figure 3.**
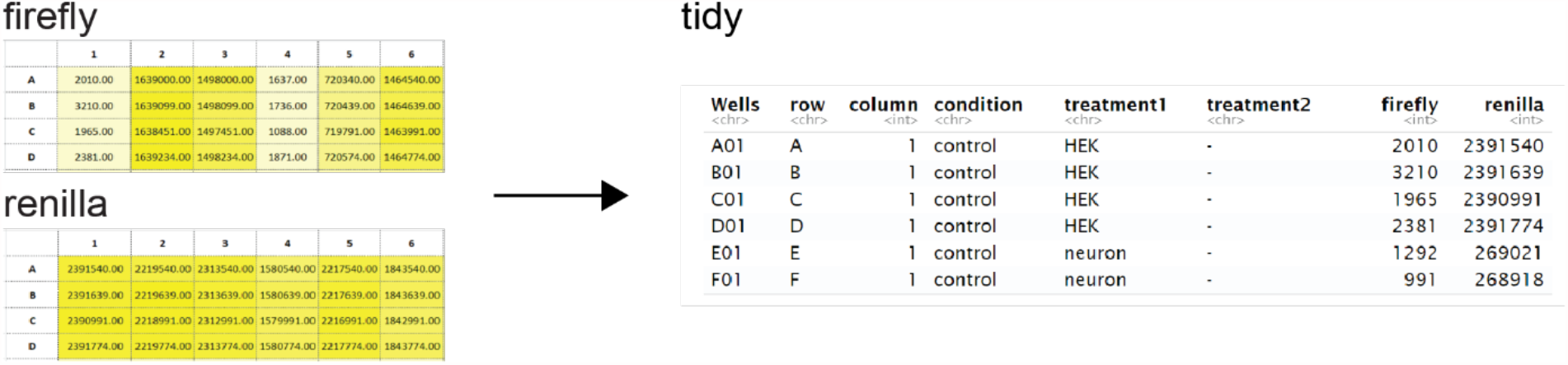
GloMax data conversion to tidy format. PlotXpress allows uploading of GloMax spreadsheet input. An additional experimental design file in tidy format should be provided to connect the reads with the conditions. After supplying the design file, the GloMax data is converted to a table in tidy format, where each row contains a single observation and the experimental conditions. The table can be downloaded and used as input for plotXpress.

#### Tidy data upload

Instead of uploading a GloMax spreadsheet in 96-well format and an additional design file, users can choose to upload a single tidy dataset (.CSV) containing both experimental conditions and firefly and renilla signals. An example file is available: https://github.com/ScienceParkStudyGroup/PlotXpress/blob/master/plotXpress_Tidy-3.csv

All the relevant data from one experiment (design and measurements) should be present in one tidy table. Users are free in the size of the experiment: any number of cell types, sequences of interest, or treatments may be added. The minimal information that is required is a column with wells (in the format A01, B01, ..), a column with intensity data and a column with the conditions. An additional column with reference data is optional and is used to normalize the data if supplied.

### Data handling

In dual luciferase assays, each replicate consists of a firefly luciferase and renilla luciferase intensity reading. To normalize the reporter gene expression to the internal control, plotXpress calculates the ratio of firefly signal to renilla signal, resulting in the firefly/renilla ratio. An average firefly/renilla ratio is then calculated for each group of readings that share identical names in the “condition” column. Finally, the fold change is calculated by dividing the firefly/renilla ratio by the firefly/renilla ration of a selected reference condition:

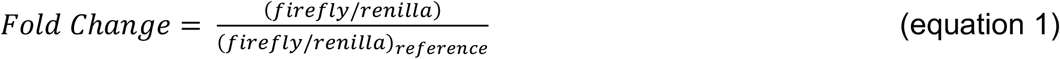

PlotXpress offers flexibility in the selection of conditions to filter the data before plotting. Individual wells can be removed from the analysis or specific conditions can be filtered.

The combined dataset including information about the wells, condition, treatments, firefly and renilla signals and the calculated ratios can be downloaded in tidy format (CSV) for further processing and statistical testing.

## Data visualization

In the plotting tab called “Plot”, a reference condition can be selected. Without selecting a reference, the firefly/renilla ratio is plotted on the y-axis. If a reference condition is selected, normalized firefly/renilla values from the other conditions are represented as fold-change relative to the reference (**Fig 4A**).

**Figure 4.**
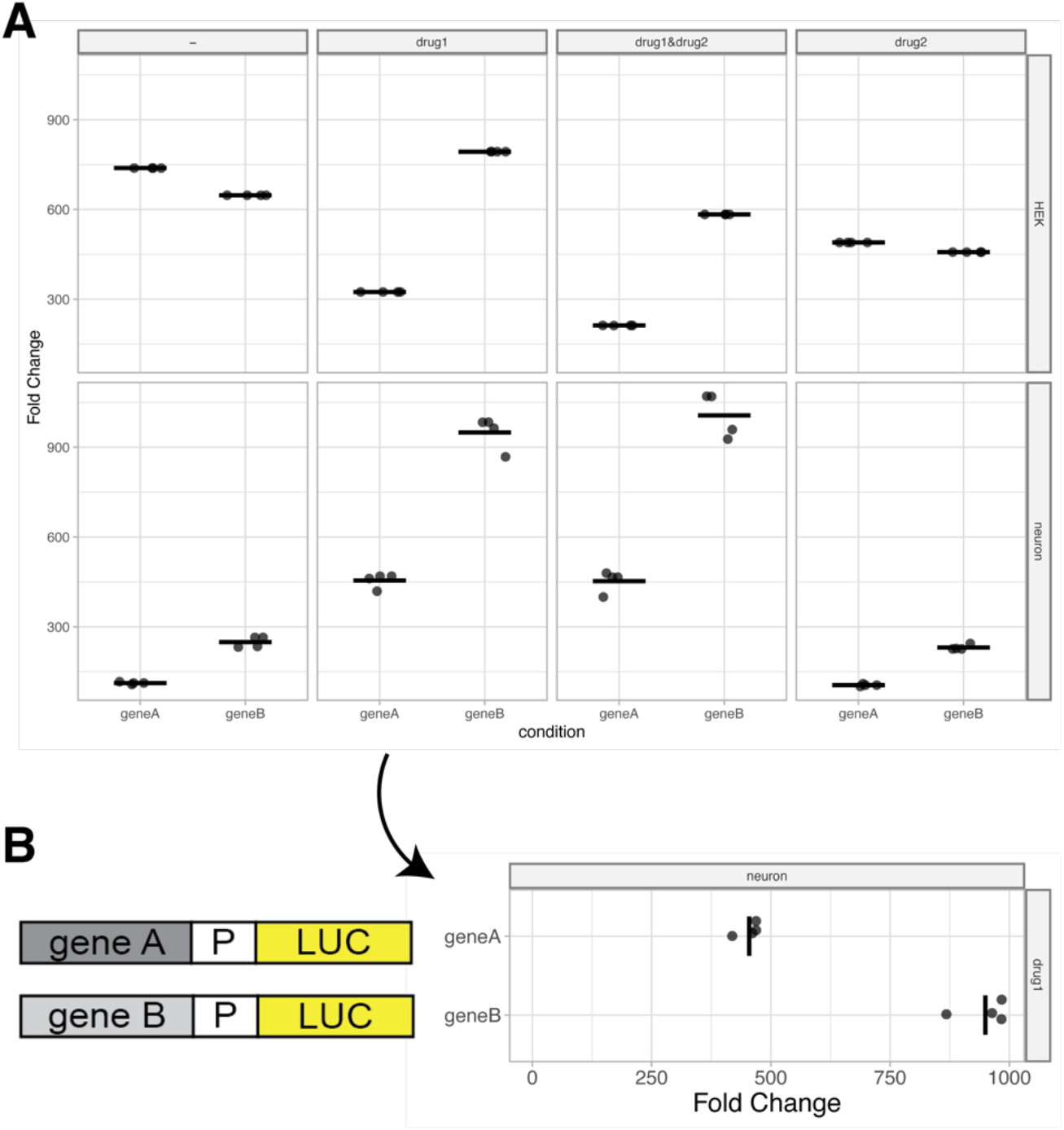
Plots based on example data. (A) Standard output plot produced by plotXpress after selecting a reference condition. The plots show the fold change in expression induced by gene A and gene B relative to a reference, for each of 8 different conditions. Individual datapoints are shown and the median of each condition is indicated with a horizontal line. (B) A 90° rotated panel for one of the conditions (drug1 in neurons) shown in panel A. Rotation of the panel improves readability of the fold change and allows for the display of a cartoon of the reporter gene next to it.

In plotXpress a categorical condition can be compared by selecting an option in the “compare” section. This condition is shown on the x-axis. Each datapoint of the reporter assay is plotted as a dot and represents the normalized firefly/renilla ratio. Dot size and transparency can be adjusted as well as other features such as axis scales, plot title, font size and the use of grid lines. The mean or median can be shown per condition as a line (Fig 4A). When presenting reporter expression data, it can be useful to display a rotated plot. Diagrams of reporter constructs can be included to visually support the experimental set up and show the corresponding measurements in line (**Fig 4B**). As an optional feature, the reference can be shown or hidden, depending on user preference.

## Outputs

The combined data from an experiment can be downloaded as a CSV file in tidy format. This data format can be used for further processing, such as statistical testing and plotting. The tidy format is well handled by statistical software (R) and other web tools that we have developed (available at https://huygens.science.uva.nl/). For instance, when the data consists of a mix of technical and biological replicates, this can be visualized as a superplot (Lord et al., 2020) by uploading the data in the SuperPlotsOfData app (Goedhart, 2020). Final plots can be downloaded from PlotXpress in PNG or PDF format, the latter being ideal for downstream processing with vector-based graphic software.

PlotXpress also produces a summary table in multiple formats such as CSV or PDF that can be included as supplemental data in publications (**Fig 5**)

**Figure 5.**
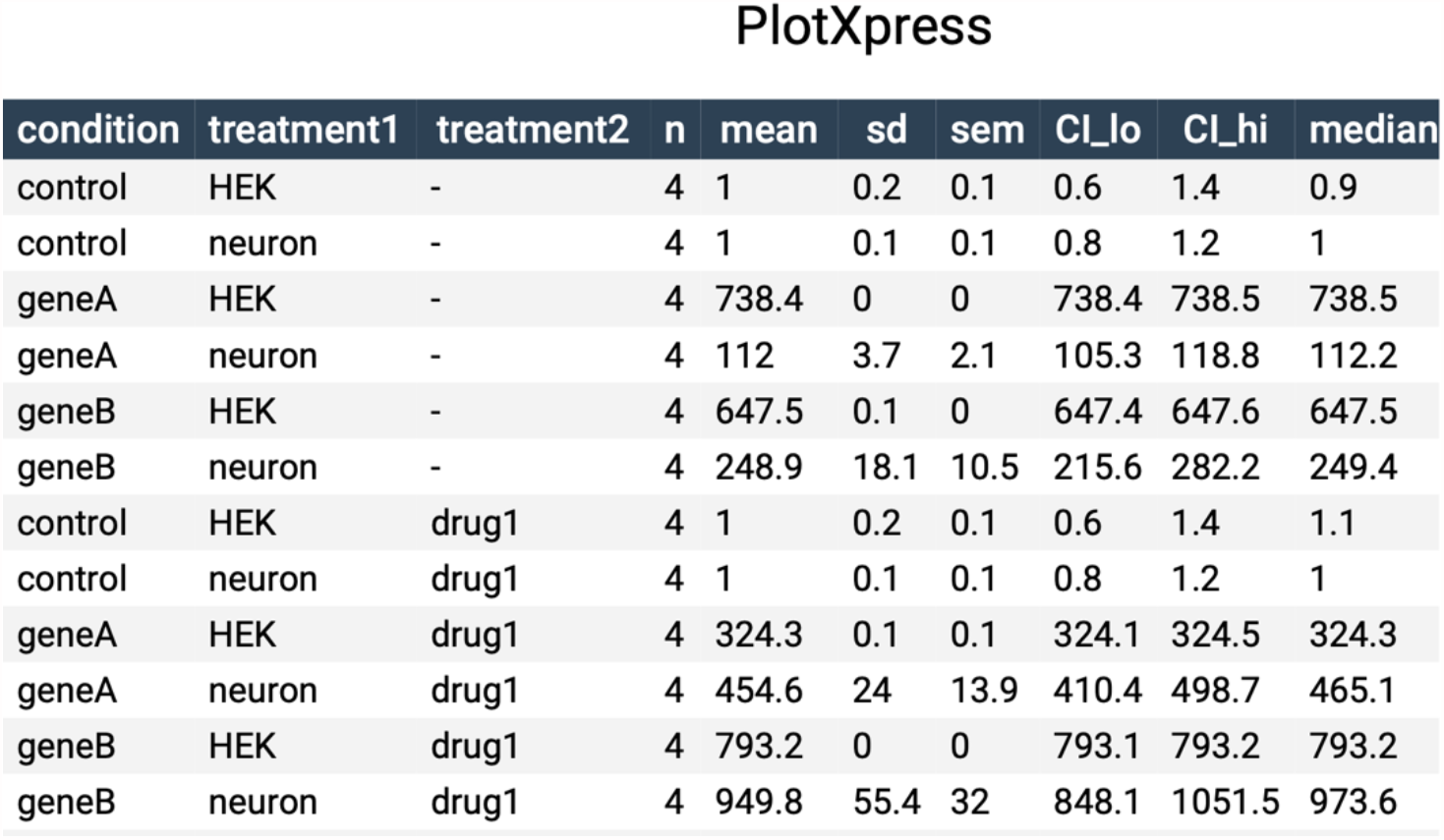
Data summary table. Table containing summary statistics of the data shown in figure 4A.

## Perspectives

The current version of plotXpress was written to analyze expression data in tidy format as well as for GloMax output data. The import of data produced by plate readers from other manufacturers is not supported at this moment, but we welcome suggestions and example data to implement import functions for other data formats. Since plotXpress is open-source, users can modify the source code by making a fork to the GitHub repository and create a pull request to have their changes reviewed and possibly integrated. Although plotXpress was developed for the analysis and visualization of dual-luciferase experiments, it can be used for other types of data that require normalization or comparison with a reference condition, such as qPCR data.

## Conclusion

Gene reporter assays are an invaluable tool for molecular biology, enabling the study of regulatory DNA sequences, transcription factors, pharmaceutical compounds, oligonucleotides and other factors that interact with DNA sequences. With each treatment or cellular context, the assay increases in complexity. PlotXpress simplifies the data analysis by bridging the gap between wet-lab standards (96-well format) and dry-lab conventions (tidy data format) that are typically used in downstream analysis of experimental biological data. The open-source web tool enables anyone to benefit from standardized data processing and state-of-the-art data visualization.

## Data availability

The data and code are deposited at zenodo: https://doi.org/10.5281/zenodo.4980068

## Author contributions

E.B. initiated the project, contributed experimental data, wrote code and wrote the manuscript. M.G. supervised the project, wrote code, reviewed code and edited the manuscript. J.G. wrote code and edited the manuscript.

## Acknowledgements

We thank our colleagues at the Swammerdam Institute for Life Sciences, University of Amsterdam, for their interest and support. We thank dr. F.M.J. Jacobs for providing resources and accommodation for this project.

